# All driven by energy demand? Integrative comparison of metabolism of *Enterococcus faecalis* wildtype and a glutamine synthase mutant

**DOI:** 10.1101/2021.07.07.451427

**Authors:** Seyed Babak Loghmani, Eric Zitzow, Gene Ching-Chiek Koh, Andreas Ulmer, Nadine Veith, Ruth Großeholz, Madlen Rossnagel, Maren Loesch, Ruedi Aebersold, Bernd Kreikemeyer, Tomas Fiedler, Ursula Kummer

## Abstract

Lactic acid bacteria (LAB) play a significant role in biotechnology, e.g. food industry, but also in human health. Many LAB genera have developed a multidrug resistance in the past few years, becoming a serious problem in controlling hospital germs all around the world. *Enterococcus faecalis* accounts for a large part of the human infections caused by LABs. Therefore, studying its adaptive metabolism under various environmental conditions is particularly important. In this study, we investigated the effect of glutamine auxotrophy (Δ*glnA* mutant) on metabolic and proteomic adaptations of *E. faecalis* in response to a changing pH in its environment. Changing pH values are part of its natural environment in the human body, but also play a role in food industry. We compared the results to those of the wildtype. Our integrative method, using a genome-scale metabolic model, constrained by metabolic and proteomic data allows us to understand the bigger picture of adaptation strategies in this bacterium. The study showed that energy demand is the decisive factor in adapting to a new environmental pH. The energy demand of the mutant was higher at all conditions. It has been reported that Δ*glnA* mutants of bacteria are energetically less effective. With the aid of our data and model we are able to explain this phenomenon as a consequence of a failure to regulate glutamine uptake and the costs for the import of glutamine and the export of ammonium. Methodologically, it became apparent that taking into account the non-specificity of amino acid transporters is important for reproducing metabolic changes with genome-scale models since it affects energy balance.

## Introduction

Lactic acid bacteria (LAB) are gram-positive microorganisms, fermenting hexose sugars to lactic acid as their primary product under many conditions. Among LABs there are both pathogenic as well as commensal species. In some cases, e.g. in the case of *Enterococcus faecalis* (*E. faecalis*) both commensal, as well as pathogenic behavior occurs^1^. As a part of the commensal flora, *E. faecalis* colonizes different tracts in the human body, especially the gut. Due to its pathogenic potential, *E. faecalis* frequently causes nosocomial infections, most commonly of the urinary tract, but also soft tissue or intra-abdominal infections, bacteremia or endocarditis^2^. An increasing proportion of *E. faecalis* strains isolated from such infection shows multidrug resistance against a wide range of antibiotics^3,4^. The treatment of infections caused by these multi-resistant *E. faecalis* strains can be remarkably hard and may cause severe problems in hospital environments^5^.

On the other hand, *E. faecalis* strains generally regarded as safe (GRAS) are used in food industry as a cheese starter culture^6^ or as a probiotic^7^. The intended probiotic isolates, however, should undergo screening to ensure the absence of transferable virulence factors and antibiotic resistant genes^8^. Hence, *E. faecalis* encounters very different native environments ranging from different human body tissues to different kinds of food. This requires enormous flexibility of the metabolism of *E. faecalis* that in turn would be reflected by various metabolic phenotypes.

To gain a comprehensive understanding of metabolic phenotypes, the cell-wide and integrative analysis of metabolism is central. This is something that pure experimental research cannot deliver, and therefore, different computational approaches have been developed to study metabolic networks. For smaller networks and more detailed analysis, kinetic models based on ordinary differential equations (ODEs) are the best choice^9^. When integrating all reactions of the metabolic network and in the absence of detailed kinetic data, genome-scale metabolic models, as used below, are nowadays the preferred and most commonly used strategy. Genome-scale metabolic models are stoichiometric representations of all annotated metabolic reactions in a given cell which allow the computation of flux distributions based on the knowledge of their localization, wiring and the biomass composition of the specific organism and/or cell type^10^. Optimal flux distributions are calculated according to an optimality criterion like biomass maximization^11^. This is an especially successful criterion for investigating microorganisms since these often follow relatively simple principles like optimizing growth. However, the typical outcome of such an optimization (flux balance analysis (FBA)) is not a unique solution and the huge size of the solution space renders interpretation of the results difficult and error-prone^12^. By adding constraints, e.g. through experimentally measured medium composition, input and output fluxes of metabolites^13^, transcriptome^14^, as well as proteome data^15^ the solution space can effectively be decreased and the predictive power of the models increased^15,16^.

In this study, we analyzed the metabolic and proteomic profile of a knock-out mutant of glutamine synthetase (Δ*glnA)* of the multiresistant *E. faecalis* V583 strain during a pH shift experiment. Glutamine Synthetase (GlnA) is a vital protein as it is the main enzyme in the assimilation of ammonia and has an overall control over the nitrogen metabolism^17^. We designed an experiment to investigate the effect of glutamine auxotrophy on the metabolic behavior of the organism under two pH conditions by comparing the results to those of the wildtype^18^. For this purpose, a previously published genome-scale metabolic model^19^ of the wildtype was adjusted to represent the Δ*glnA* mutant. Experimental data of the Δ*glnA* mutant was then used to constrain the solution space of the model. The results were compared with a likewise constrained wildtype model, thereby providing an integrative view on the metabolic adjustments that the organism has to perform to react to the imposed glutamine auxotrophy during environmental pH changes.

## Materials and Methods

### Experimental

#### Bacterial strains and culture conditions

*Enterococcus faecalis* V583 Δ*glnA*^19^ mutant was grown in batch cultures at 37 °C in a chemically defined medium for lactic acid bacteria (CDM-LAB^13^, pH 7.5 and 6.5). The CDM-LAB medium contained the following per liter: 1 g K_2_HPO_4_, 5 g KH_2_PO_4_, NaHCO_3_, 0.6 g ammonium citrate, 1 g acetate, 0.25 g tyrosine,0.24 g alanine, 0.5 g arginine, 0.42 g aspartic acid, 0.13 g cysteine, 0.5 g glutamic acid, 0.15 g histidine, 0.21 g isoleucine, 0.475 g leucine, 0.44 g lysine, 0.275 g phenylalanine, 0.675 g proline, 0.34 g serine, 0.225 g threonine, 0.05 g tryptophan, 0.325 g valine, 0.175 g glycine, 0.125 g methionine, 0.1 g asparagine, 0.2 g glutamine, 10 g glucose, 0.5 g L-ascorbic acid, 35 mg adenine sulfate, 27 mg guanine, 22 mg uracil, 50 mg cystine, 50 mg xanthine, 2.5 mg D-biotin, 1 mg vitamin B12, 1 mg riboflavin, 5 mg pyridoxamine-HCl, 10 mg p-aminobenzoicacid, 1 mg pantothenate, 5 mg inosine, 1 mg nicotinic acid, 5 mg orotic acid, 2 mg pyridoxine, 1 mg thiamine, 2.5 mg lipoic acid, 5 mg thymidine, 200 mg MgCl_2_, 50 mg CaCl_2_, 16 mg MnCl_2_, 3 mg FeCl_3_, 5 mg FeCl_2_, 5 mg ZnSO_4_, 2.5 mg CoSO_4_, 2.5 mg CuSO_4_, and 2.5 mg (NH4)_6_Mo_7_O_24_.

#### pH shift experiments in chemostat cultures

The pH shift experiments were carried out as previously described^18^. In short, *E. faecalis* V583 Δ*glnA* was grown in glucose-limited chemostat cultures in Biostat B Plus benchtop bioreactors (Sartorius) in 750 ml CDM-LAB with a dilution rate of 0.15/h at 37 °C and gassing with 0.05 L/min nitrogen and stirring with 250 rpm. The pH was kept at the desired level by titrating with 2 M KOH. Initially, the pH was kept constant at 7.5 until a steady state was reached. Steady state was assumed when no glucose was detectable in the culture supernatant anymore and dry mass and optical density (600 nm) were constant on two consecutive days. For the pH shift, the pH control was switched off until the desired pH (6.5) value was reached. The cultivation was continued until the steady-state was reached again. Samples were taken at steady state pH 7.5 and at several time points during and after the pH shift as indicated in Figure 1. Per sampling point, samples for determination of dry mass, extracellular metabolites and proteomic analysis were taken as previously described^18^.

**Figure 1.**
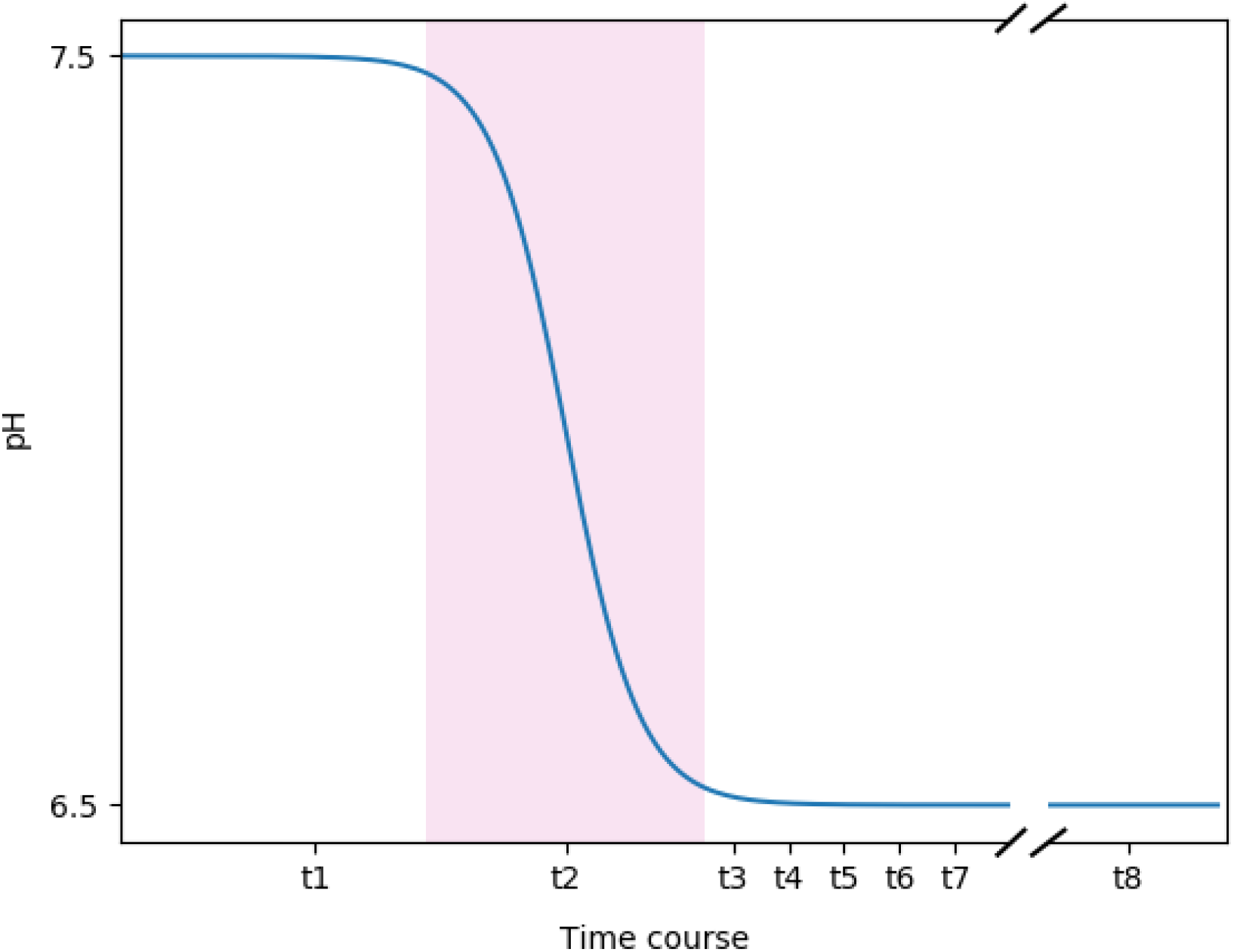
Time course of the pH shift experiment. Samples were taken at t1 (steady-state at pH 7.5), t2 (the transition state (pink background) during the pH shift), t3, t4, t5, t6 and t7 which indicate data points at pH 6.5 at 80, 100, 120, 180 and 240 minutes after the start of the pH shift. Finally, samples were taken at t8, the steady-state at pH 6.5, 21 h after the start of the experiment. The distance between the data points does not represent the actual time difference in the experiment. Also, the break between t7 and t8 shows the shortened x axis between the two data points.

#### Chemostat cultures for determination of ATP_maintenance_

For determination of ATP_maintenance_ (ATPm), *Enterococcus faecalis* V583 Δ*glnA* was grown in glucose-limited chemostats as described above (except for pH shift) at two different dilution rates, 0.15 h^−1^ and 0.05 h^−1^, with three biological replicates per dilution rate. At steady-state samples were taken and processed as described above.

#### Quantification of extracellular metabolites

For samples from pH shift experiments, quantification of amino acids in media and culture supernatants was done by Frank Gutjahr Chromotgraphie (Balingen, Germany) and quantification of lactate, formate, acetate, glucose, acetoin, 2,3-butanediol, ascorbate, citrate, pyruvate and ethanol were done by Metabolomics Discoveries GmbH (Potsdam, Germany). For quantification of amino acids, glucose, and fermentation products in CDM-LAB and culture supernatants of samples from ATP_maintanance_ experiments, the following two methods were used:

Method 1: an Agilent 1260 Infinity II HPLC system was used. The system was controlled by OpenLAB CDS Workstation software. For the amino acids analysis, sample supernatants were filtered through a 0.22 μm syringe filter into a HPLC sample vial. Amino acids were derivatized, separated on a reversed-phase column (Agilent Poroshell 120 EC-C18 4.6×100mm, 2.7μm), detected with a diode array detector (DAD G7117A) and quantified following manufacturer’s guidelines (AdvanceBio Amino Acid Analysis, © Agilent Technologies, Inc. 2018). Standards ranging from 5 μM to 30 mM were used for the quantification of aspartate, glutamate, asparagine, serine, glutamine, histidine, glycine, threonine, arginine, alanine, tyrosine, valine, methionine, tryptophan, phenylalanine, isoleucine, leucine, lysine and proline.

For analysis of organic compounds, samples were prepared as follows: 100 μl 35 % perchloric acid were added to 1 ml sample, mixed and placed on melting ice for 10 minutes. Subsequently, 55 μl potassium hydroxide solution (7 M) were added and the sample was centrifuged for 2 min at 20,000 *g*. The supernatant was filtered through a 0.22 μm syringe filter into a HPLC sample vial. Separation of sugars and fermentation products in the sample was performed by using an Agilent Hi-Plex H column (4.6×250 mm, 8 μm) with a working temperature of 65 °C using 10 mM H_2_SO_4_ as a mobile phase with a flow rate of 0.4 ml/min. For detection, a refraction index detector (RID) with a working temperature of 35 °C and a diode array detector (DAD) with a wavelength of 210nm/4nm with a reference wavelength of 360nm/100nm were used. Standards ranging from 50 μM to 150 mM were used for the quantification of glucose, ethanol, citrate, lactate, pyruvate, formate and acetate.

Method 2: Sugars and organic acids in the supernatant were measured with an isocratic Agilent 1200 series HPLC system equipped with a Phenomenex guard carbo-H column (4 by 3.0 mm) and a Rezex ROA organic acid H (8%) column (300 by 7.8 mm, 8 μm; Phenomenex) maintained at 50°C. Analytes were separated and detected using 5 mM H2SO4 with a constant flow rate of 0.4 mL min^−1^. Prior to analysis samples were pretreated for precipitation of abundant phosphate by addition of 4 M NH3 and 1.2 M MgSO4 solution followed by incubation with 0.1 M H2SO4. Absolute concentrations were obtained by standard-based external calibration and normalization with L-rhamnose as internal standard.

#### Quantification of protein abundances

All the steps in quantification of protein abundances were exactly carried out as previously described in Großeholz *et al*^18^.

### Computational

#### Determination of non-growth associated ATP_maintenance_

The determination of non-growth associated ATP (ATPm) was performed as described in Teusink et. al^20^. Thus, the measured flux value for the carbohydrates, organic acids and amino acids were integrated in the genome-scale model as constraints. The biomass reaction was fixed at the respective growth rate (dilution rate) and the flux of the ATPm reaction was maximized as the objective function. The obtained values were used to fit a linear function, for which the *y*-intercept determines the required energy for the organism at zero growth rate. This value is then applied to the model as the lower bound of the ATPm reaction.

#### Integration of constraints to the genome-scale model

The integration of constrains to the genome-scale model were done as indicated in Großeholz *et al*^18^. To integrate the metabolic data a tolerance level of 40% was applied to the measured flux rates to account for measurement errors. The obtained values were applied to the upper (+ 20%) and lower (−20%) bounds of the respective exchange reactions at both conditions. Regarding the proteome data, the reaction with no experimental evidence at the proteome level at pH 7.5 were deactivated. To represent the significant fold changes of proteins in response to pH shift, the log2 change of protein abundances were multiplied by 40% (tolerance level) and then applied to the maximum and minimum value of respective reactions, obtained by flux variability analysis (FVA)^21^ at pH 7.5.

## Results

In order to follow the metabolic adjustments to pH changes in the Δ*glnA* mutant of *E. faecalis* V583 a chemostat set-up was used and a pH shift from pH 7.5 to 6.5 was applied. The respective pH profile can be seen in Fig. 1. At all indicated time-points, samples were taken and subjected to biomass, metabolite and proteome measurements. The results are compared with earlier measurements of the corresponding *E. faecalis* wildtype strain under the same conditions^18^.

### Effect of pH on the growth rate

The biomass production of *E. faecalis* Δ*glnA* decreased from 1.54 to 1.15 g/l when the pH was shifted from 7.5 to 6.5 (Fig. 2). The trend of decreasing the biomass production at lower pH values is similar to the wildtype, as the biomass production in both genotypes decreased by approximately 25% in response to pH shift. However, the biomass production of the wildtype at any given pH value is larger than that of the mutant, suggesting an important role of the glutamine synthetase reaction for the growth of the organism.

**Figure 2.**
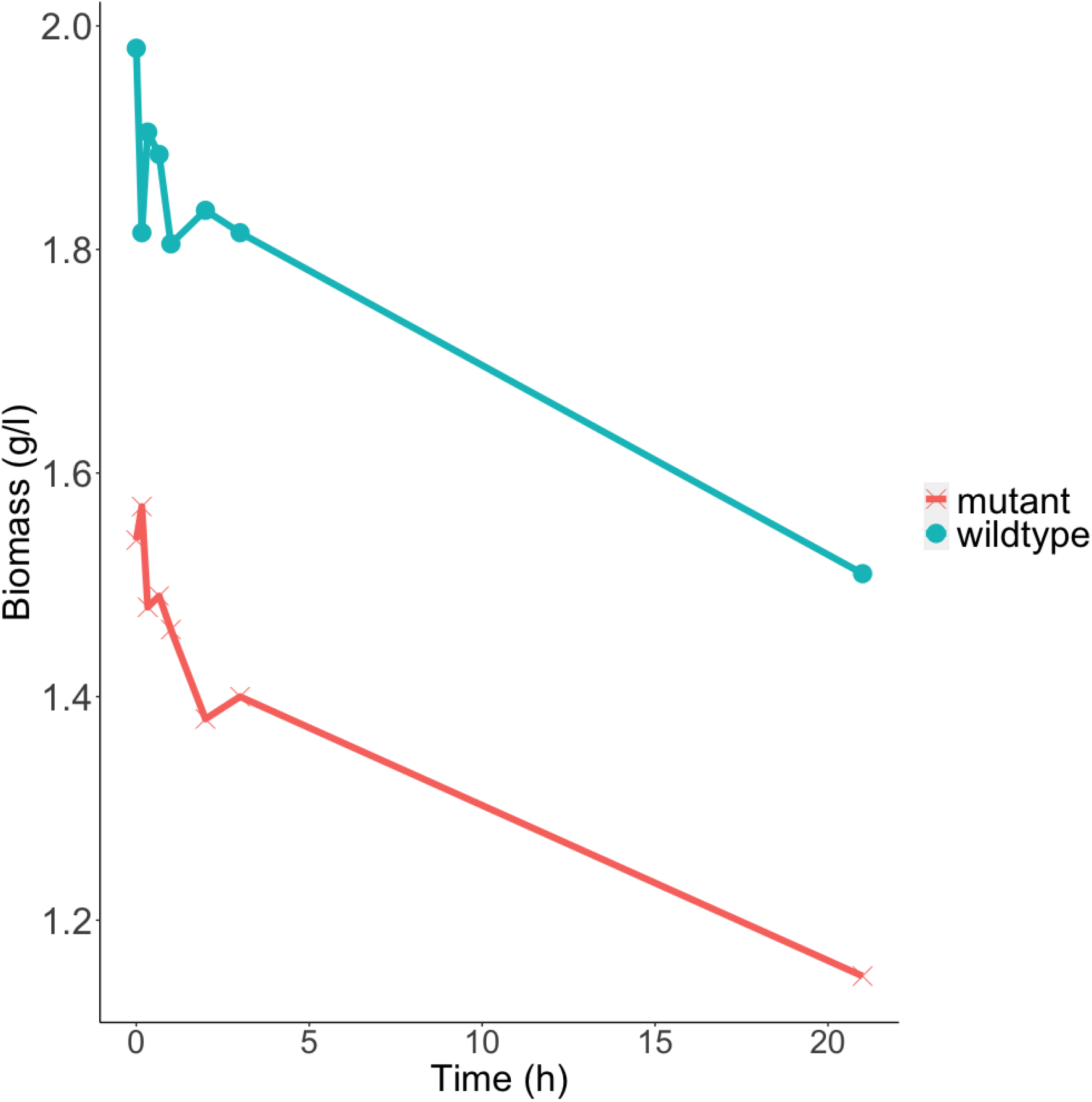
Development of the biomass production over the course of 21 hours during the pH shift experiment. The red line shows the biomass values of the E. faecalis Δglna mutant, while the blue line shows the one of the wildtype. Each data point represents the mean of two biological replicates.

### The effect of pH on metabolite uptake and production

To discover the effect of a pH shift on the metabolic behavior of *E. faecalis* Δ*glnA,* the concentration of the extracellular carbon source (glucose), organic acids as well as amino acids was determined in the samples of the chemostat experiment and the respective uptake or production rates were calculated accordingly (Fig. 3). The measured profile of the carbon source and organic acids consists of the uptake rate of glucose, as the primary energy source for the organism, and the fermentation profile containing lactate, acetate, ethanol and formate, reflecting the state of the energy metabolism at each pH value. Similar to the wildtype, the glucose uptake rate increased in response to the drop in pH, indicating a higher energy demand in a more acidic environment. This is mostly caused by the need to pump protons out of the cell at the expense of ATP^18^. Moreover, and also similar to the wildtype, the fermentation pattern changed from mixed acid fermentation to homolactic fermentation, as homolactic fermentation is energetically more favorable. Despite the qualitative similarity of the pH response to that of the wildtype, quantitatively, the uptake rate of glucose in the mutant showed a stronger increase compared to the wildtype. This is translated to a higher lactate production, suggesting a higher energy demand in response to pH shift in the Δ*glnA* mutant. Also, the extent of shift to homolactic fermentation is stronger in the mutant. So, in summary, these results indicate a higher energy demand in the Δ*glnA* mutant of *E. faecalis* compared to the wildtype at all pH values.

**Figure 3.**
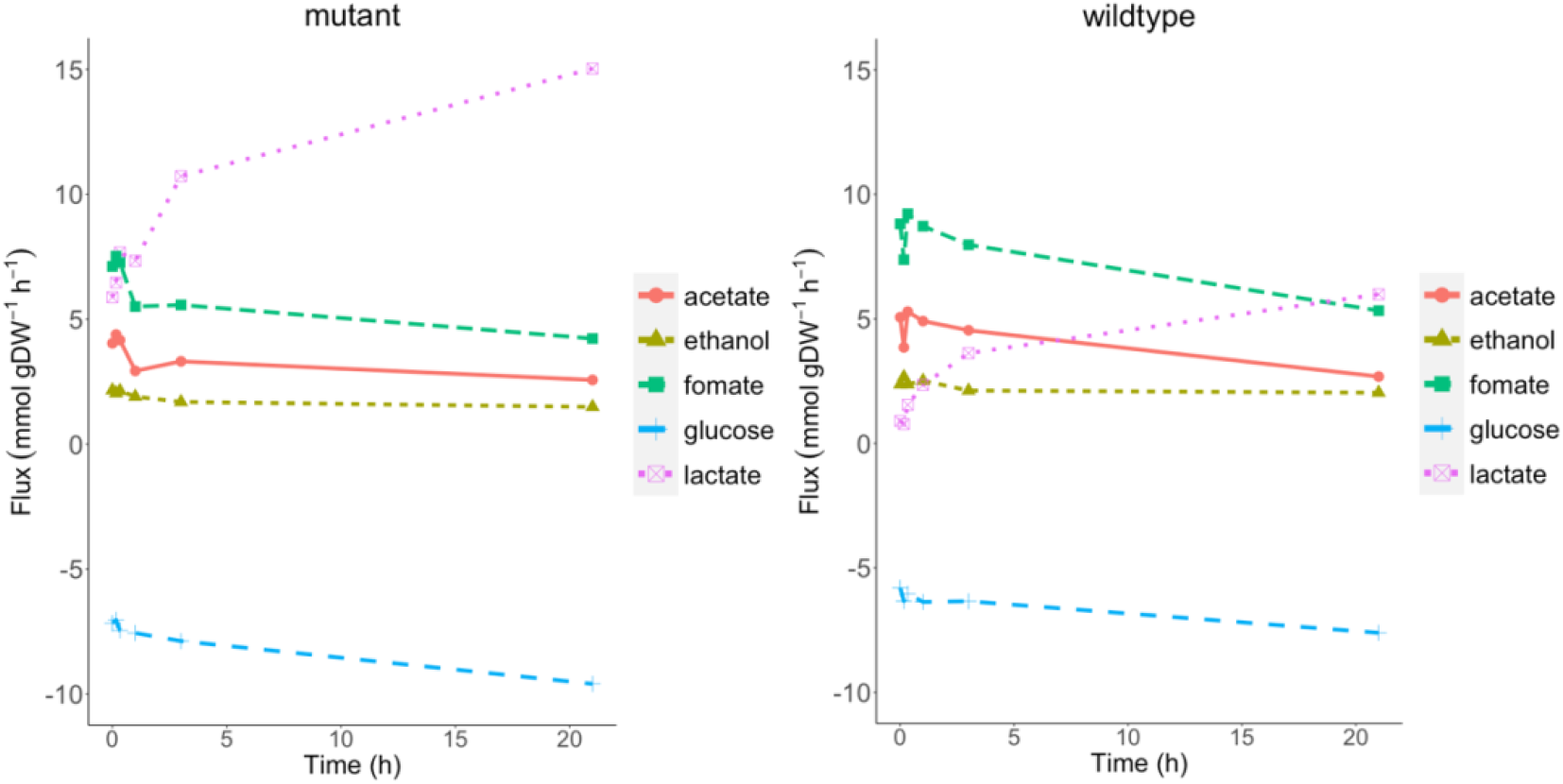
The uptake and production rate of glucose and fermentation products during the pH shift experiment. The left panel shows the data from the mutant, and the right panel shows the data from the wildtype. Each data point represents the mean of two biological replicates.

This increased energy demand is also suggested when comparing the uptake and production rates of amino acids between *E. faecalis* Δ*glnA* and the wildtype (Fig. 4). Overall, the amino acids uptake rate decreased in response to pH shift with the exception of arginine, glutamine and serine. Since less biomass is produced and more ATP is needed for proton export, less protein synthesis should occur. At the same time, amino acid uptake is also energy consuming since it is accompanied by either direct ATP consumption or additional proton import. Therefore, it is interesting to look at the reason for an increased uptake rate under these conditions. The uptake rate of arginine and serine, as well as the production rate of ornithine both increased in the wildtype and the mutant after the pH shift. It has been reported that the catabolism of arginine via arginine deaminase is used by a variety of lactic acid bacteria in response to a more acidic environment^22^. Initially, it had been believed that there is a beneficial buffering by ammonia. But when calculating the actual stoichiometries, we previously showed that this is not the case at the respective pH^13^. However, arginine is readily metabolized to gain ATP which can be used to pump protons to the extracellular environment under a more acidic condition. Therefore, it can be suggested that under more energy-demanding conditions (mutant versus wildtype, and pH 6.5 versus 7.5 (in both genotypes)), a higher uptake rate of arginine may help cells to boost energy production. Serine uptake was also increased after the pH shift in the Δ*glnA* mutant, as serine can as well be used for ATP production via degradation to ammonia and pyruvate and fermenting pyruvate to acetate.

**Figure 4.**
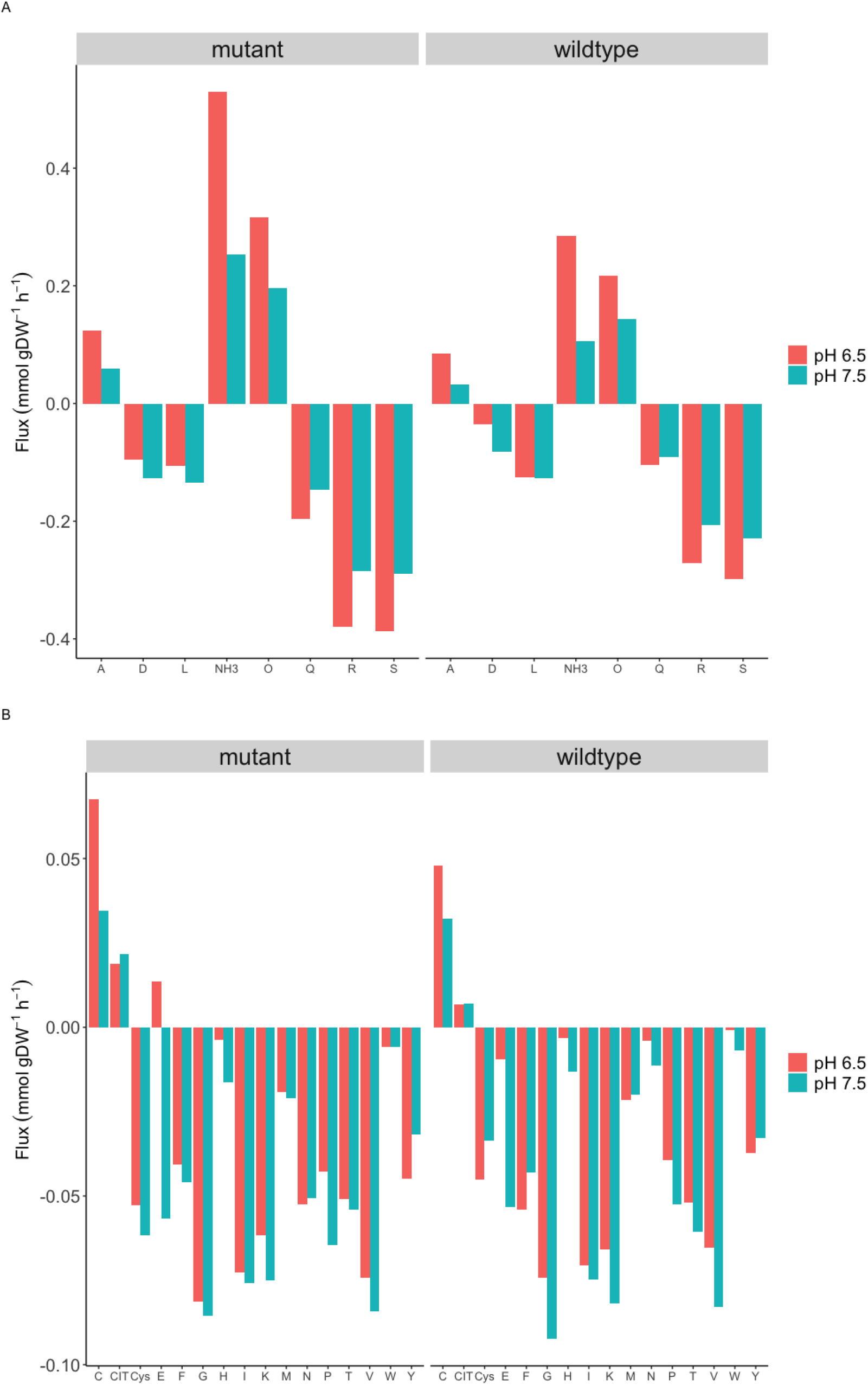
The uptake and production of the amino acids in the wildtype and the ΔglnA mutant at pH 7.5 and 6.5 (t1 and t8, respectively). Panel A shows the uptake/production rate of amino acids with high flux value (larger than 0.1 mmol/g^−1^_DW_h^−1^) and panel B shows the uptake/production rate of amino acids with low flux value (smaller than 0. mmol/g^−1^_DW_h^−1^)

Additionally, the uptake rate of glutamine and the production rate of ammonia were considerably higher in the mutant compared to the wildtype at both pH levels. The latter is a curious observation, since the export of such amounts of ammonia points to excess nitrogen from glutamine. The glutamine auxotrophy of the mutant might explain the larger margin between the two pH values, as the higher uptake rate ensures that the growth is less affected by the auxotrophy. More importantly, however, it has been reported that in *Streptococcus pneumoniae*^23^ the transcription factor GlnR (which controls the production and transport of glutamine) is dependent on the intact gene for GlnA to successfully function. The overproduction of ammonia suggests that the regulatory effect of GlnR is also disrupted in *E. faecalis* Δ*glnA*, resulting in the upregulation of the glutamine ABC transporter (GLNabc) and an unnecessarily high glutamine uptake rate accordingly. This also accounts for the small glutamate excretion in the mutant at pH 6.5, since massive amounts of glutamine will drive the glutamine deaminase to produce both glutamate and ammonium. The glutamate excretion was however not observed in a previous study^19^.

### Significant fold changes on protein level during the pH shift

To observe the effect of the pH shift on the protein expression in *E. faecalis* Δ*glnA,* the expression rate of all detected proteins was quantified throughout the pH shift experiment and the respective significant fold changes were calculated at different time points compared to time point 1 (t1). The complete set of significantly changed protein expressions is shown in the supplement (supplementary table 1). While there was no significant fold change at t2, the highest number of fold changes was observed at t3, 20 minutes after the start of the pH shift, with more than 40 proteins (out of 1681 detected ORFs) being affected. Among all affected proteins eleven are involved in membrane and cell wall production, and two proteins assigned to reactions involved in peptidoglycan biosynthesis. This suggests that restructuring of the membrane and cell envelope occurs early in response to a change in environmental pH - similar to what has been observed for the wildtype^18^. A smaller number of proteins was affected by the pH shift between t4 (1 hour after start of the pH shift) and t7 (4 hours after start of the pH shift), all of which were downregulated. At t8, 21 hours after the pH shift, the number of significant fold changes amounted to 40, with the majority of the proteins being downregulated. A large number of these is involved in nucleotide biosynthesis (Table 1). Considering the fact that *de novo* biosynthesis of nucleotides is an energy demanding process for the organism, the down regulation of the respective pathways is in line with the higher energy demand under the more acidic condition.

**Table 1.** The number of significant changes in protein abundances and the prominent respective subsystems at t8 (ΔglnA mutant)

| Subsystems | Number of affected genes |
| --- | --- |
| Amino acid metabolism | 3 |
| Carbohydrate metabolism | 7 |
| Nucleotide metabolism | 22 |
| Transport | 6 |

To find out the differences between the wildtype and the mutant at the proteome level in response to pH shift, the significant fold changes were compared. Except for t8, the number of significant fold changes in the wildtype was considerably higher than that in the mutant. As already mentioned, at early time-points, both the mutant and the wildtype showed fold changes in enzymes that are involved in membrane and cell wall production. However, at these early time-points after pH shift the wildtype also displayed an upregulation of glycolytic enzymes compared to the mutant. The lack of increased expression in glycolytic enzymes in the mutant might be explained by the previously introduced higher energy demand. Since glycolysis is responsible for the major energy production in *E. faecalis*, it is plausible that the expression of those enzymes was already at a higher level in the mutant. The downregulation of enzymes involved in nucleotide metabolism after the pH-shift is again similar between wildtype and mutant.

### Computational

In order to study the metabolic behavior of *E. faecalis* Δ*glnA* more comprehensively and on a cell-wide scale, genome-scale modelling in combination with constraint-based modelling was applied. For that matter, the previously published genome-scale metabolic network of *E. faecalis*^19^ was used to simulate the pH shift experiment by the integration of metabolic and proteomic data from the *E. faecalis* Δ*glnA*. To integrate the experimental data into the genome-scale model, the framework from our previous^18^ work was applied. To represent the *glnA* knock-out in the model, the reaction flux of glutamine synthetase was set to zero. The complete model is available at biomodels.

### Determination of the non-growth associated ATP value of *E. faecalis* Δ*glnA*

In order to prepare the genome-scale model of *E. faecalis* Δ*glnA* for the integration of the above described data and the analysis via FBA, we determined the non-growth associated energy demand (ATPm) which is an important feature for performing FBAs. The non-growth associated energy demand reflects the amount of energy required to sustain life at zero growth. Thus, the definition of this parameter is an important aspect for genome-scale modelling as it can strongly affect the flux distributions in the metabolic network.

For this purpose, *E. faecalis* Δ*glnA* was grown in chemostat cultures at two different dilution rates (0.05, 0.15 h^−1^). After reaching the steady-state, samples were taken from the chemostat cultures and the metabolite composition in the supernatant was experimentally determined to calculate the uptake/production rate of extracellular metabolites (Table 2). The uptake and production rates of glucose, organic acids, and amino acids were then integrated into the model as reaction constraints, defined as bounds on the respective exchange reactions. To calculate parameters for the ATPm reaction, for each dilution rate, the biomass production reaction was fixed at the maximal growth rate that is equal to the value of the dilution rate, and the flux through the ATPm reaction was maximized as the new objective function. The obtained maximal objective function value, namely the maximal flux through the ATPm reaction at different dilution rates is plotted against the respective dilution rate and a linear function was fitted to those values. The non-growth associated energy reflects the amount of energy that microbes require to survive at zero growth and is derived from the *y*-intercept of the linear function. This value represents the maintenance energy and was used as the lower bound of the ATPm reaction in the model.

This constraint ensures that the minimum required amount of energy for non-growth associated purposes is produced by the model and is not channeled into biomass production and thus into growth. As a result, the calculated value of ATPm of the *E. faecalis* Δ*glnA* was 5.977 and 6.224 mmol/g^−1^_DW_h^−1^ at pH 7.5 and 6.5, respectively. When integrated into the model, however, the maximal biomass production is too high compared to experimental data. In fact, the level at which the model produced the fermentation products resulted in a very high ATP production, so at this value of ATPm, the ATP is redirected into biomass production. Hence, the constraints of this reaction had to be set on a higher value under both pH conditions. Several reasons allow for such adjustment without violating the modeling rules. First, the ATPm is an estimated value by taking the measured value of around 30 metabolites into account. Therefore, the estimation is very error prone, as the measurement error of all the metabolites accumulate and impact the optimization process. For instance, the glucose uptake rate at pH 7.5 at the dilution rate of 0.15 was 6.32 mmol/g^−1^_DW_h^−1^ in the data set used for ATPm estimation, and 7.17 mmol/g^−1^_DW_h^−1^ in another previously measured data set shown in Figure 3. This difference of 0.85 mmol/g^−1^_DW_h^−1^ results in approximately 2 mmol/g^−1^_DW_h^−1^ difference at the optimized ATPm value at this particular condition. Second, the ATPm value in essence is meant to ensure that the model is not reaching the optimal growth by overlooking the non-growth associated energy. In this sense, deviation from the estimated value to a higher value does not violate the underlying assumption, while deviation to a lower value would require more solid evidence. Here, both calculated values for the ATPm reaction seem to be too low and had to be increased. If we consider lactate production as an indicator of the energetic state of the cell, the fact that the lactate production in the mutant is several times higher than in the wildtype suggests a much higher ATPm value for the mutant compared to the wildtype (the ATPm values in the wildtype were calculated to be 3.9 mmol/g^−1^_DW_h^−1^ at pH 7.5 and 8.4 mmol/g^−1^_DW_h^−1^ at pH 6.5^18^). Therefore, the ATPm values in the mutant were increased to 9.7 mmol/g^−1^_DW_h^−1^ and 10.6 mmol/g^−1^_DW_h^−1^ at pH 7.5 and 6.5, respectively. For pH 7.5, this represents the minimal value for the correct reproduction of biomass production. For pH 6.5 we selected a higher value, but noted that the exact value does not considerably impact the solution. These values allowed for a precise prediction of the biomass and the production rates of the organic acids. It is necessary to point out that under both pH conditions, in addition to being glucose limited, the chemostat cultures were also deprived of glutamine, suggesting that the glutamine content of the CDM-LAB might be limiting and might thus be insufficient to fulfill the demands in *E. faecalis* Δ*glnA.*

### Predicted flux through the energy metabolism in the Δ*glnA* mutant

The experimentally measured metabolic and proteomic data were integrated into the *ΔglnA* genome-scale model to simulate the pH shift experiment (as described for the wildtype^18^). Thus, in short, the uptake and release fluxes are set as boundaries for the respective fluxes and fluxes associated with proteins not present in the proteome data are set to zero, if not essential. Finally, adjustments of fluxbounds according to de- or increases in expression are implemented.

As reflected by the experimental data, the *ΔglnA* mutant has a higher energy demand at all conditions. This obviously also holds true in the model after the integration of experimental data. The model shows an increased flux through glycolysis, which implies a higher ATP production. Accordingly, the predicted flux through lactate dehydrogenase (LDH) is also increased after the pH shift.

The flux distribution in glycolysis was compared between wildtype and mutant to investigate the difference in flux values in energy metabolism. At pH 7.5 a higher flux value passed through the glycolytic reactions in the mutant as a result of the higher energy demand. In the model, the higher energy demand in the mutant results from the need to import glutamine from the extracellular environment either via the GLNabc (glutamine ATP binding cassette) transporter or via glutamine permease. GLNabc-mediated uptake of glutamine consumes ATP and permease-mediated uptake imports protons into the cell, which subsequently increases the ATP demand as ATP is required to pump protons out of the cell. However, glutamine synthesis from glutamate would of course also consume one ATP per glutamine. Therefore, the striking difference in energy demand is still not explained. We will come back to this point below.

### Impact of glutamine uptake in the model of *E. faecalis* Δ*glnA*

As explained above, the fact that the Δ*glnA* mutant is unable to produce glutamine leads to increased uptake of extracellular glutamine. Based on the chemostat data, glutamine uptake increased from t1 at pH 7.5 to time point 8 at pH 6.5, corresponding to a 34% increase from 0.147 mmol/g^−1^_DW_h^−1^ to 0.197 mmol/g^−1^_DW_h^−1^, respectively. Originally, the glutamine transport via the glutamine ABC transporter (R_GLNabc) and glutamine permease (R_Glnt6) were represented in the model. As proteome data suggested, the ATP binding subunit of the GLNabc (EF0760) was upregulated after the pH shift, suggesting an increase of the transporter demand which is consistent with the higher uptake rate of glutamine. Considering the fact that membrane proteins were often missing from the proteomic data set (due to technical issues), it is likely that the other subunits of the transporter were subjected to upregulation as well, as there is an increase in glutamine uptake after the pH shift. However, the original design of the glutamine transport in the model could not translate the upregulation of GLNabc into a higher flux. The FBA flux distribution revealed a flux value of zero for GLNabc under both pH conditions. In order to gain a consistent result between the model and the experimental data, an improved, new permease reaction was introduced to the model. As reported previously, the permease system in *E. faecalis* is not amino acid-specific, but is rather shared between multiple amino acids with various affinities^24^. It is suggested that while glutamine has the highest affinity to the transporter, the same transporter takes up asparagine and threonine as well. Hence, to account for the higher affinity of the transporter for glutamine compared to the other amino acids, the new permease reaction in the model was designed to carry two glutamines together with one asparagine and one threonine and four protons (one per each amino acid molecule). The previous amino acid specific permease of all three amino acids were then deactivated accordingly. The new transport design resulted in a successful prediction of the uptake rate of all three amino acids and also in using GLNabc when a higher uptake rate is in demand after the pH shift. The new set up accounts for the actual transport system in a more accurate way, as it mimics the shared permease system and also gets the ABC transporter in use when needed. The fact that during glutamine uptake involuntarily and automatically other amino acids are taken up as well (expending ATP or taking up protons) might be also at least partially the source for the high energy demand of the mutant. In addition, and as mentioned above, the regulatory effect of GlnR in *E. faecalis* Δ*glnA* might be disrupted so that the uptake of glutamine (and the other amino acids transported by the same proteins) is less strictly regulated.

### Predicted flux through glutamine/glutamate metabolism

The flux distribution of the genome-scale model was subsequently used to analyze the effect of the pH shift on glutamine-glutamate metabolism. This pathway is of particular interest as we aimed to uncover the consequences of glutamine auxotrophy in the metabolism of *E. faecalis* in this study. Following an increase in glutamine uptake and on the contrary, decrease in glutamate uptake, the model predicted an upregulation in the glutamine to glutamate conversion, which was reflected in switching on the two reactions, aspartyl-tRNA(Asn):L-glutamine amido-ligase (ADP-forming) (ASNTAL) and carbamoyl-phosphate synthase (glutamine-hydrolysing) (CBPS). Interestingly, the model predicted a flux shutdown at glutamine-fructose-6-phosphate (gam6p) transaminase, which produces glucose amine-6-phosphate by a transaminase reaction between glutamine and fructose-6-phosphate. Instead, gam6p is produced by assimilating ammonia into fructose-6-phosphate. The model also predicted an increased flux towards the reverse direction of glutamate dehydrogenase (GDH), producing glutamate from 2-oxoglutarate. The directionality of GDH plays an important role in balancing the carbon and nitrogen metabolism^25^. The NADPH/NADP ratio is directly influenced by less NADPH being available for e.g. amino acid biosynthesis when the flux is directed towards glutamate production. This also coincides with our observation of a strongly decreased uptake of glutamate from the medium and an increased uptake of glutamine (which is required for 2-oxoglutarate production). The decreased level of NADPH also prevents reductive synthesis reactions from taking place, as it is reflected in the significant downregulation of proteins involved in e.g. nucleotide metabolism. As another beneficial side effect, the reverse direction of the GDH consumes one proton. This flux change also leads to a series of changes in other amino acid production/degradation processes. For instance, a higher conversion rate of glutamate to alanine and aspartate was predicted, with the former being excreted by the cell after the pH shift, based on the experimental data.

## Discussion

*E. faecalis* is important in the food industry and hospital environments; therefore, characterizing its highly adaptive metabolism is necessary. Hence, Integrative analysis of bacterial metabolism is a key approach to uncover their adaptive behavior. In this study we analyzed the effect of a decline in environmental pH from 7.5 to 6.5 on a *ΔglnA* mutant of *E. faecalis* and compared the results to those of the wildtype^18^. Like the wildtype, the *ΔglnA* mutant responded to the pH shift by reprogramming its metabolic and proteomic profile to fit the increased energy demand that comes with the need to maintain a higher pH by pumping protons out of the cell. Many findings therefore paralleled the results from the wildtype thereby also confirming these. However, there are also striking differences, mostly concerning the quantity of the energy demand in the genotypes.

Similar to the wildtype, the mutant decreased biomass production during pH shift since less ATP is available for anabolic processes. In addition, there is a pronounced shift from mixed acid to homolactic fermentation. While the qualitative pattern in the mutant resembled the one in the wildtype, the proportion of lactate production in the mutant was considerably higher. This confirms that under more energy-demanding conditions (*ΔglnA* mutant or acidic conditions), *E. faecalis* changes its fermentation profile to homolactic fermentation. While the stoichiometric analysis of the fermentation pathway shows that mixed acid fermentation produces one more ATP, it is widely reported that LAB species such as *L. lactis* and *L. plantarum* choose homolactic fermentation during high glycolytic flux, high substrate availability or faster growth rates^26^. The high uptake rate of glucose under energetically demanding conditions, which further translates into a higher glycolytic flux, increases the NADH/NAD ratio. Reportedly, a higher ratio of NADH/NAD upregulates the activity of lactate dehydrogenase (LDH), e.g. in *L. lactis*^27^. While this has been observed in the literature, a plausible explanation why the less productive (in terms of ATP) fermentation is favored under energetically demanding conditions is still missing.

The amino acid uptake/production profile further underlines the overwhelming influence of the growing energy demand when pH is lowered. The uptake rate of amino acids mostly decreased following the pH shift except for arginine, serine and glutamine. A decreased amino acid uptake should be a result of regulation when less biomass is produced, and also saves energy, since the uptake is either coupled to even more proton uptake or to ATP hydrolysis. The increased uptake of arginine and serine can be easily explained since these can be directly used for ATP production. Arginine is considered a prolific energy resource in many lactic acid bacteria, especially those important in the food industry^28,29^. An interesting aspect of arginine catabolism in lactic acid bacteria (when used for energy production) is that it often does not result in citrulline production, regardless of using arginine deaminase or not^28,29^. Likewise, our data suggested that an increase in arginine uptake following the pH shift leads to an increase in ornithine, but not citrulline production. The previous finding on the production of ornithine from citrulline through ornithine carbamoyl transferase in *E. faecalis*^30^ was supported by the flux distribution in the genome-scale model. Ornithine is then used by the arginine-ornithine antiporter to import even more arginine into the cell.

Not surprisingly, there is quite a big difference between wildtype and mutant in glutamine/glutamate metabolism and how this is affected by the pH shift. While the uptake rate of glutamate considerably decreased in response to pH shift, the glutamine uptake rate was increased – in the wildtype only very slightly and strongly in the mutant. As discussed above, several mechanisms might contribute to this observation – first, we proposed in accordance with literature on *S. pneumoniae*^23^ that the regulatory effect of the transcription factor GlnR on the uptake of glutamine depends on the intact gene for GlnA and its absence results in an unregulated glutamine uptake. Second, the reversal of the flux of the GDH with its regulatory consequences is only possible, if more 2-oxoglutarate is produced for which glutamine is needed. Third, as learned from the genome-scale model, it is important to consider the lack of specificity of the amino acid uptake mechanisms. Co-transported with e.g. glutamate are amino acids like aspartate that cannot readily be catabolized for energy production. In the case of glutamine, at least small amounts of arginine are also co-transported which is favorable under high energy demand.

The glutamine synthetase reaction (catalyzed by GlnA) is the main reaction to assimilate ammonium^17^. However, here glutamine is imported in such large quantities that some of it has to be deaminated to produce glutamate (so that even a small amount of excretion is observed) and ammonium which the model also predicted. Although high concentrations of ammonium are reported to lower the growth rate in bacteria, the underlying reason is suggested to be the general osmotic or ionic effect of ammonium rather than its toxicity^31^. The exact mechanism of ammonium export is not known, but likely either ask for proton antiport or ATP which adds to the energy demand of the mutant.

The analysis of the proteome data again showed a lot of parallel adjustments to pH shift between wildtype and mutant. Here, the decrease of expression in enzymes of the *de novo* biosynthesis of nucleotides which is costly, and also the upregulation of enzymes involved in the restructuring of the cell membrane and cell wall which is necessary while facing a drop in extracellular pH level in order to decrease proton leak is common between the genotypes. The mutant however shows a striking lack of increasing the protein expression of glycolytic enzymes at the beginning of the pH shift experiment. Since there is already a high glycolytic flux in the mutant at the start of the experiment, we assume that the respective changes in core metabolism already happened at this point.

All experimental data were reproducible in the genome-scale model. However, the stoichiometry of amino acid uptake reactions had to be adjusted to a realistic depiction of their non-specificity. To the best of our knowledge, this has not considered in other genome-scale models of bacteria so far.

Initially, when considering all of the above findings which mostly reflect the higher need for energy in the mutant, it was not obvious why this higher need arises. Glutamine is the primary nitrogen donor in bacterial cells. High levels of glutamine have to be maintained in order to allow effective transfer of amino-groups^32^. This can be accomplished by its synthesis and uptake in the wildtype or its uptake alone in the mutant. At first glance the energetic cost of glutamine uptake is comparable compared to its biosynthesis via GlnA. One ATP is needed for GlnA and the usage of one ATP or proton import is the consequence of glutamine import. However, if the control over glutamine uptake is inhibited due to the lack of GlnA, the potentially uncontrolled and excessive import of glutamine may incur additional cost to the cells. Thus, both the automatic co-transport of unwanted amino acids, as well as the need to excrete large amounts of ammonia are certainly costly.

**Figure 5.**
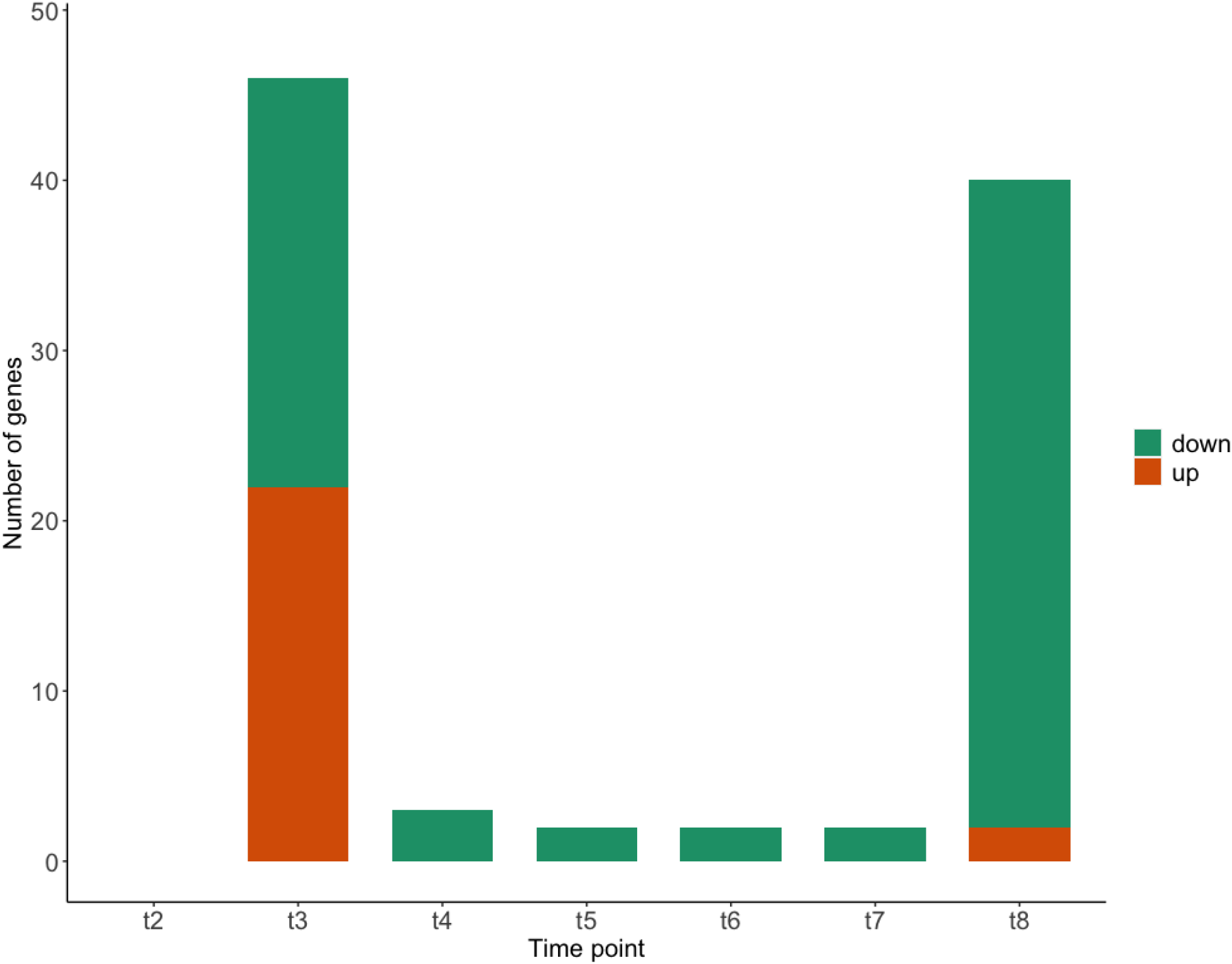
The significant change in protein abundances in the ΔglnA mutant during the pH shift experiment at different time points.

**Figure 6.**
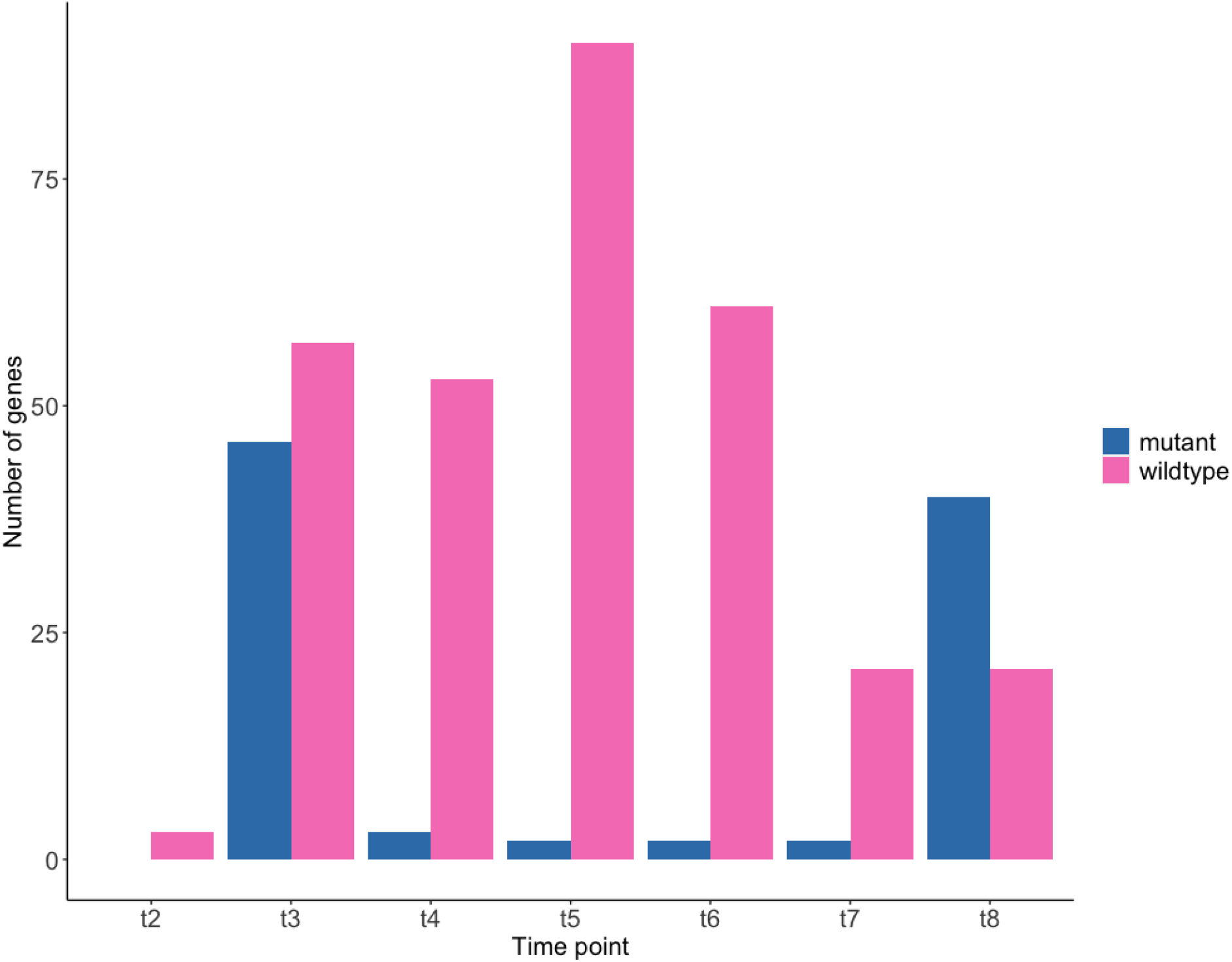
The number of significant changes in protein abundances during the pH shift experiment in the wildtype and the ΔglnA mutant.

## Acknowledgements

SBL would like to thank the HGS Mathcomp and the Graduate academy at Heidelberg University for financial support. NV and TF thank the DFG (grant numbers VE1075/2-1 and FI 1588/ 2-1) for funding. Ben C Collins and Olga T Schubert, the co-authors of Großeholz *et al* are acknowledged for proteomics analysis.

